# Improving the analysis of biological ensembles through extended similarity measures

**DOI:** 10.1101/2021.08.08.455555

**Authors:** Liwei Chang, Alberto Perez, Ramón Alain Miranda-Quintana

## Abstract

We present new algorithms to classify structural ensembles of macromolecules, based on the recently proposed extended similarity measures. Molecular Dynamics provides a wealth of structural information on systems of biologically interest. As computer power increases we capture larger ensembles and larger conformational transitions between states. Typically, structural clustering provides the statistical mechanics treatment of the system to identify relevant biological states. The key advantage of our approach is that the newly introduced extended similiarity indices reduce the computational complexity of assessing the similarity of a set of structures from O(*N*^2^) to O(*N*). Here we take advantage of this favorable cost to develop several highly efficient techniques, including a linear-scaling algorithm to determine the medoid of a set (which we effectively use to select the most representative structure of a cluster). Moreover, we use our extended similarity indices as a linkage criterion in a novel hierarchical agglomerative clustering algorithm. We apply these new metrics to analyze the ensembles of several systems of biological interest such as folding and binding of macromolecules (peptide,protein,DNA -protein). In particular, we design a new workflow that is capable of identifying the most important conformations contributing to the protein folding process. We show excellent performance in the resulting clusters (surpassing traditional linkage criteria), along with faster performance and an efficient cost-function to identify when to merge clusters.

## Introduction

Computational Biology approaches complement our experimental knowledge to understand Biological phenomena at different timescales and length scales, and helps tame the combinatorics of possible sequences leading to functional proteins, peptides or nucleic acids. Molecular dynamics-based techniques capture the physics of these macromolecules at the atomic scale, following their time evolution at the femtosecond regime. At this timescale, they provide the mechanistic details that might be too fast for current experiments to capture.^1^

Thanks to improved supercomputer architecture and software development, these tools can now routinely simulate events well into the microseconds,^2, 3^ bringing insight into a plethora of biological processes. Events that require longer timescales, such as binding or folding, rely on advanced sampling techniques that preserve the thermodynamics and in some cases the kinetics as well.^4–9^ Simulations produce ensembles of structures which require a statistical mechanics treatment to identify relevant states of interest from the computational results.

Most often, we are interested in identifying the metastable states in the system, and the microstates that belong to them. Traditionally, this is done by using some similarity measure between conformations (often a 2D-RMSD calculation).^10^ An alternative approach relies on projecting trajectories onto a set of features, and then use some dimensionality reduction approach (PCA or tICA).^11^ In both cases, a clustering method classifies structures into different states—with the largest population state corresponding to the lowest free energy basin. In the case of folding or binding predictions, the centroid of the most populated cluster is compared against experimental structures. However, consideration of larger time- and length-scales inevitably means an explosive growth in the number of structures/conformations that must be analysed. This means that developing more efficient data analysis techniques is paramount to study more realistic biological systems.

Here we propose a novel take on this problem, by applying our recently developed extended similarity indices^12, 13^ to the study of conformational ensembles. These indices generalize many of the most popular similarity measures commonly used in cheminformatic studies, but they have a key difference: while standard indices can only quantify the similarity between two objects (e.g., they are binary functions),^14, 15^ our indices can calculate the similarity between an arbitrary number of objects at the same time (e.g., they are *n*-ary functions). A key advantage of this approach is that we can perform the similarity calculations with unprecedented efficiency: calculating the similarity of *N* objects using binary indices scales as O(*N*^2^), while the *n*-ary indices scale as O(*N*). Moreover, the truly global nature of our extended indices provides a more natural description of the similarity of a group of objects, which results in better estimates of set compactness. We recently showed how to compute the *n*-ary indices for sets of vectors formed by an arbitrary number of categorical characters,^16^ however, here we will apply the original formalism that takes binary vectors (bitstrings) as input. This not only leads to more efficient calculations, but also provides a more convenient way to encode the 3D structure of the molecules studies via contact maps (e.g., we can represent a conformation as a vector of 0s and 1s, where a contact between two residues can be considered a 1 and its absence a 0).

We showcase the performance of our framework in several datasets relevant to an array of computational biology problems ranging from protein folding to binding events between proteins and other macromolecules. We show the accuracy and reliability of the method as well as the agreement of predicted states with previous studies. The analysis of the protein folding landscape is particularly remarkable, since we are able to determine the most representative structures contributing to this process with minimum computational cost. Additionally, it is important to highlight that many of the tools that we develop here can have far reaching applications in data analysis. For instance, we present a linear scaling algorithm to find the medoid of a set. We show how the extended similarity indices can be used as novel linkage criteria in hierarchical clustering algorithms, with results that are consistently more robust than those obtained with other linkages. It is also noteworthy that the extended similarity-based clustering provides a natural and efficient way to estimate the number of clusters present in a dataset, an important hyperparameter that is often quite difficult to determine. Overall, our approach shows great promise at the time of analysing large and complex datasets, which can be easily generalized towards on-the-fly similarity comparisons due to the trivial cost of increasing the ensemble size.

## Methods

### Choice of Datasets

We selected datasets that were representative of a heterogeneous set of problems in computational structural biology. The datasets correspond to different ensembles extracted from molecular dynamics approaches for studying protein folding, and the binding of proteins to DNA, peptides or proteins. Each type of dataset represents unique challenges due to the heterogeneity of the ensembles. These datasets provide practical examples for the application of the clustering technology.

#### Protein-peptide datasets

The p53-MDM2 interaction has been widely studied in cancer research. P53, an intrinsically disordered protein, binds MDM2 through a small peptide epitope that adopts a helical conformation upon binding MDM2. The interaction marks p53 for degradation. Drug design efforts target MDM2 to allow free p53 to perform its role as “guardian of the genome”. The disorder-to-order transition is hard to study and predict via computational tools. Here we look at three peptides interacting with MDM2: the native p53 (PDB: 1ycr), pdiq (PDB: 3jzs) and ATSP-7041 (peptide extracted from PDB: 4n5t). The latter two have been proposed inhibitors that have intrinsically higher helical content and lead to higher binding affinities. We used ensembles produced using the MELD approach^17^ based on published binding simulations.^18^

#### Protein-protein datasets

Most proteins assemble into larger molecular complexes to carry out their functionality. We chose a homodimer (PDB: 1ns1) and heterodimer (PDB: 2mma) chosen from a pool of proteins simulated from MELDxMD binding simulations.^19^ The large interface region between the proteins as well as internal dynamics of each system pose challenges in identifying meaningful clusters.

#### Protein-DNA dataset

We use a recent protein-DNA binding dataset based on the CREB bZIP protein dimer binding a 21 base pair DNA (PDB: 1dh3). The geometry of the two long protein helices binding perpendicular to the DNA makes it challenging to cluster together structures that belong to different binding modes. The ensembles exhibits a large binding plasticity as the protein explores interactions with different base pairs along the double helix.^20^

#### Protein folding datasets

We include three datasets from explicit solvent simulations^21^ of protein folding pathways (NTL9, α3D and NuG2) as well as ensembles using implicit solvent MELDxMD simulations to fold protein G starting from extended chains. (PDB codes: 2hba, 2a3d, 1mi0 and 3gb1). In these simulation ensembles we are concerned with the internal structure of the protein, whereas the previous simulations dealt with the relative position of two different molecules.

### Featurization of the ensembles

Our similarity measures compare N-dimensional vectors of 0s and 1s representing absence or presence of the feature in a particular frame. We map our structures to feature vectors by calculating contact maps. The presence of a contact gets labelled as a 1 and its absence as a 0. We use pyemma^22^ to process the ensembles, selecting subsets of structures in the ensemble. (see Table 1). Contacts for folding simulations included all residues, while those for binding included all the residues in one subunit against all the residues in the other subunit. The only exception is the protein-DNA system, which had long helices where only a few residues where in the binding site. Thus for the protein-DNA system we chose residues 1 to 27 and 57 to 81 in the protein for contacts with all residues in the DNA.

**Table 1.** Contact map featurization of the system for similarity calculations. *indicates that only every other residue was included in the calculation. Cα: uses Cα distances between residues to calculate a contact. Heavy: uses the smallest heavy-atom distance between tow residues.

| SYSTEM | SELECTED<br>RESIDUE | CUTOFF<br>(NM) | ENSEMBLE<br>SIZE | #<br>FEATURES |
| --- | --- | --- | --- | --- |
| **NTL9 (2HBA)** | 1-39* | 1.0 (Cα) | 5000 | 380 |
| **α3D (2A3D)** | 1-73* | 1.0 (Cα) | 5000 | 1332 |
| **NUG2 (1MI0)** | 1-56* | 1.0 (Cα) | 5000 | 784 |
| **PDIQ-MDM2** | 1-85* : 86-97 | 0.7<br>(heavy) | 5000 | 516 |
| **P53-MDM2** | 1-85* : 86-100 | 0.7<br>(heavy) | 5000 | 645 |
| **ATSP7041-MDM2** | 1-85* : 86-101 | 0.7<br>(heavy) | 5000 | 688 |
| **1NS1**<br>**(HOMODIMER)** | 1-742 : 75-146 | 0.8<br>(heavy) | 3500 | 2664 |
| **2MMA**<br>**(HETERODIMER)** | 1-107 : 108-187* | 0.8<br>(heavy) | 3500 | 4280 |
| **PROTEIN G**<br>**(3GB1)** | 1-56 | 0.6<br>(heavy) | 1923 | 1540 |
| **PROTEIN-DNA**<br>**(1DH3)** | 1-27,57-81 :<br>111-152 | 0.6<br>(heavy) | 1516 | 2132 |

### Extended similarity calculations

The extended similarity indices were calculated using the fractional weight function, with minimum possible coincidence threshold in each case (e.g., for *N* datapoints, the minimum coincidence threshold is *N* mod 2). The code to perform the analyses presented here can be found at: https://github.com/PDNALab/extended-ensemble (the code to calculate the extended similarity indices is publicly available at https://github.com/ramirandaq/MultipleComparisons).

## Results and Discussion

### Identifying native structures via efficient medoid calculation

A similarity-based analysis can be very useful to determine, given a set of structures, which is the one closest to the native (e.g., bound) conformation. This can be easily seen if we consider standard binary similarity indices. For instance, if we have a pairwise similarity index *S_ij_* between any two structures *i* and *j*, then for each structure *i* we can define its group similarity, *G_i_*, as 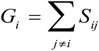. Then, we should expect that the structure with maximum group similarity is going to be the most representative of the native state. Phrased in this way, this problem is equivalent to finding the medoid of the set of structures.

The rationale behind this approach is particularly apparent in our present case, when we are representing the different protein conformations via contact maps. First of all, this provides a natural definition for *S_ij_*, which we can take as counting the number of coinciding “on” bits in the binary vectors of *i* and *j*. It is important to remark that (contrary to some similarity indices commonly used in cheminformatics) we must ignore the coincidence of 0’s. While the coincidence of 1’s means that a given contact is present in two frames, the coincidence of 0’s does not mean that two frames are similar, because in one structure the two residues could be at 10 Å and in another at 20 Å. Notice then that the bitstring of the folded state (by virtue of being compact), should have the maximum number of “on” bits. This, together with the fact that we expect to have many conformations that are close to the native structure, explains why *G_i_* should be a maximum for this conformation. We can test this hypothesis by seeing how the group similarity is correlated with the RMSD with respect to the native structure. In Fig. 1 we show the corresponding plots for two types of systems, showing that indeed *G_i_* can be used to determine the bound state. (Corresponding figures for other systems are shown in the SI.)

**Figure 1.**
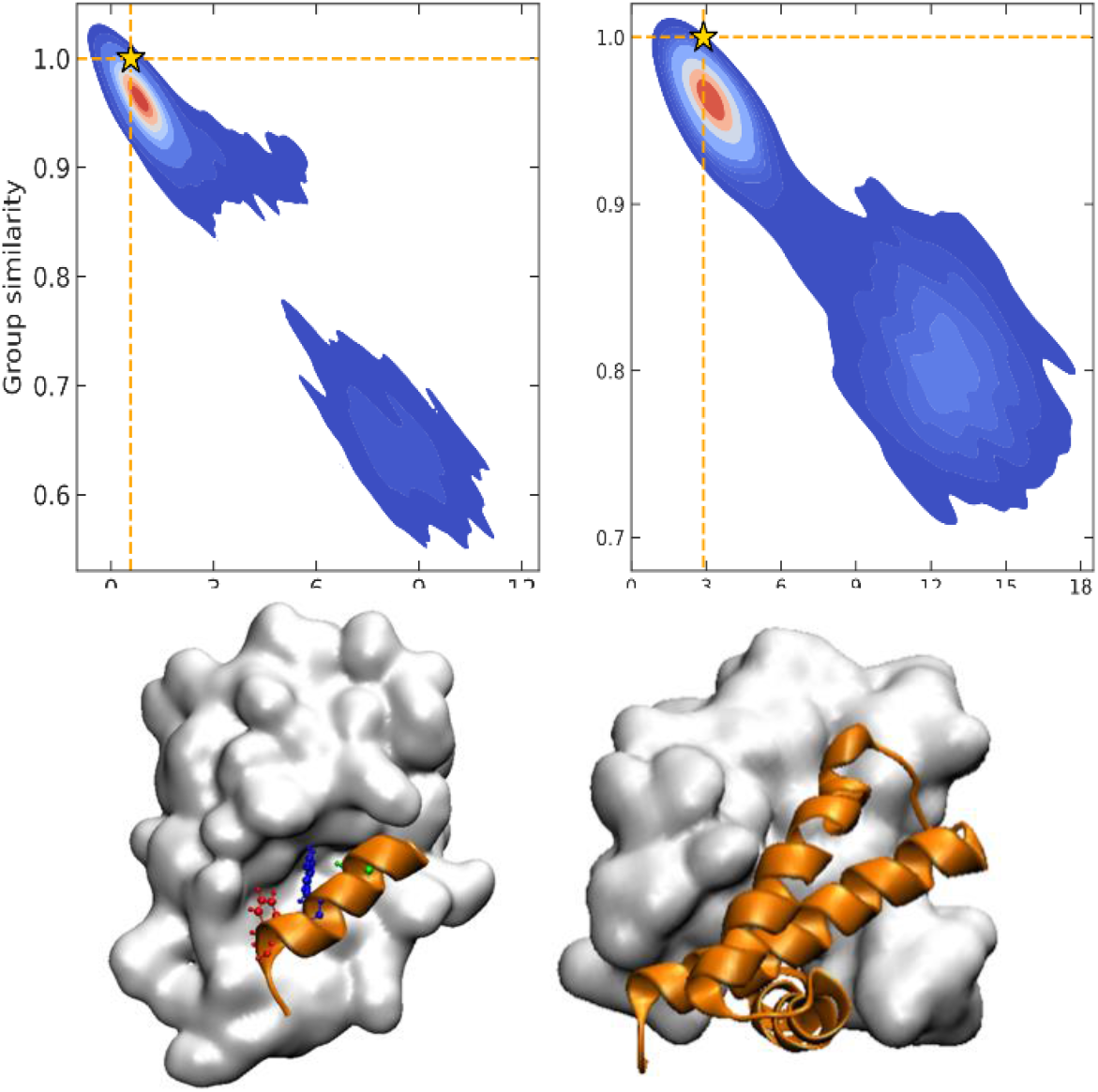
Group similarity versus RMSD against the experimental native structure for NTL9 (top left) and A3D (top right) with highest group similarity sample labeled as a star; representative structures with highest group similarity for two peptide binding systems: p53-MDM2 (bottom left) and 1NS1 (bottom right).

However, despite the promise of this approach, the fact that it hinges on binary similarities means that it has a fundamental drawback: if we have *N* structures, determining the native configuration will scale as O(*N*^2^), since we need to compute the full similarity matrix [*S_ij_*]. This is in once again in line with the medoid calculation problem, for which most algorithms scale quadratically.

An alternative route to bypass this issue would be to use *n*-ary similarity indices. For instance, given a set of *N* conformations: **C** = {*C*_1_, *C*_2_,…, *C_N_*}, we can calculate the complementary similarity of any element, 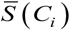, as the extended similarity, *S_e_*, of the original set without the corresponding element. That is:

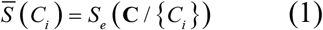

At this point we can realize that the structure with the highest group similarity also has to correspond to the one with lowest complementary similarity. The relation between these two indicators can be seen in Fig. 2, where we show how *G_i_* is almost perfectly correlated (e.g., is consistent)^23–25^ with the 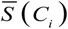 calculated using the extended (non-weighted) Russell-Rao (RR) index.

**Figure 2.**
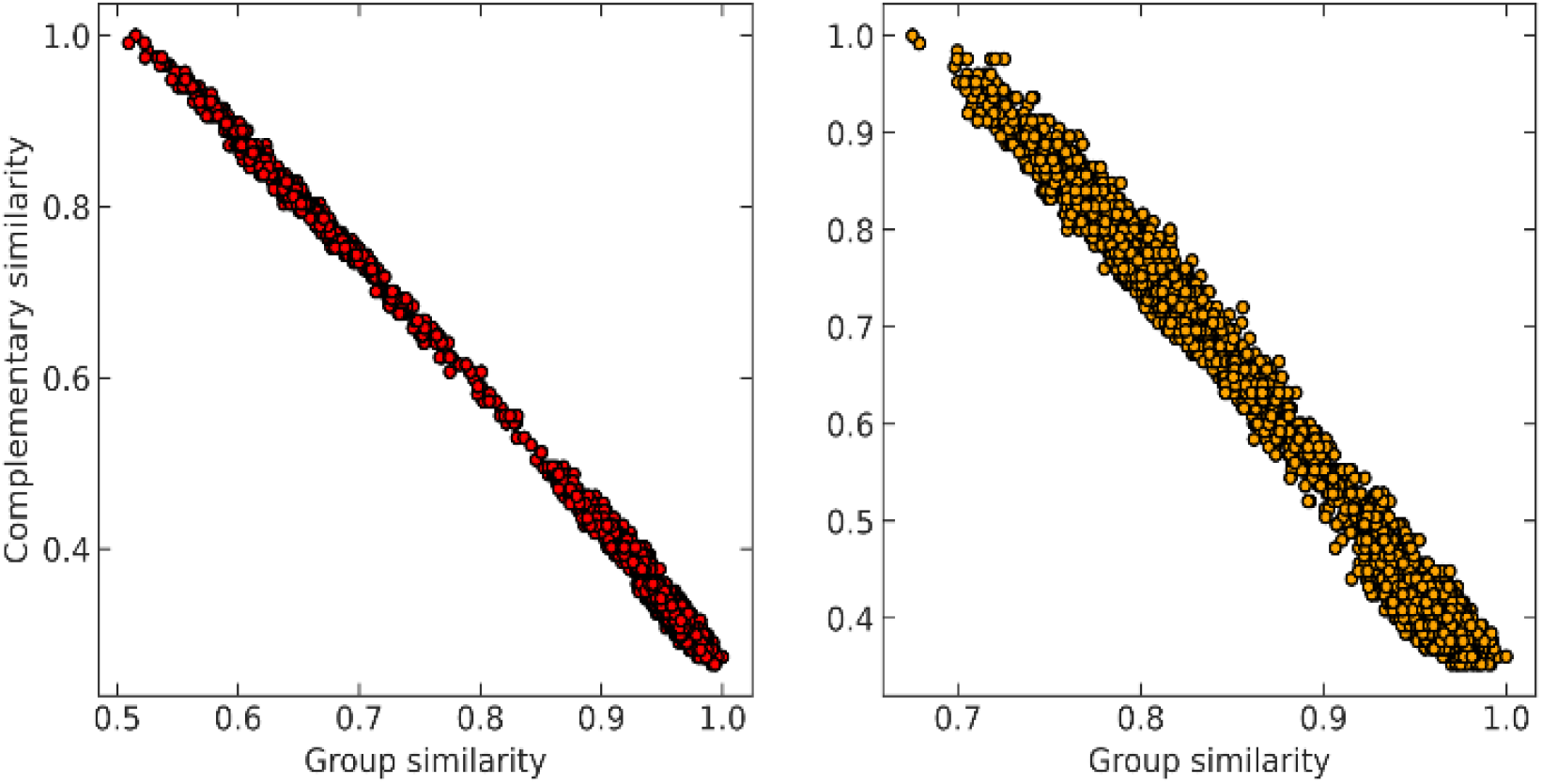
Correlation between group similarity and extended similarity using non-weighted Russell-Rao index for NTL9 (left) and A3D (right).

In other words, we can also use 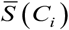 to locate the most populated state. The (very) attractive advantage of using this method, however, is that it only requires O(*N*) operations. The key insight is that given a set of bitstrings (like the contact maps that we are working with), calculating their *n*-ary similarity only requires the column-wise sum of the matrix formed by these binary vectors. With this, we can propose the following algorithm:

1. Given a set **C** = {*C*_1_, *C*_2_,…, *C_N_*}, calculate the column-wise sum of all the contact maps: ∑_C_.
2. For all conformations, subtract the corresponding bitstring from the total column-wise sum: ∑_C_ – *C_i_*.
3. Use these vectors to calculate the complementary similarity: 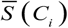.
4. Select the structure with minimum value of 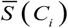.

The ∑_C_ – *C_i_* vector calculated in the 2^nd^ step is all we need to calculate any *n*-ary index over the set **C** / {*C_i_*}. Finally, note how the time-consuming steps (1^st^ and 2^nd^) scale as O(*N*), resulting in an unprecedentedly efficient (linear) algorithm to estimate the medoid of a set of binary vectors.

### Extended similarity-based clustering

The previous section deals with a very important application of the *n*-ary indices, but determining the closest structure to the folded state only requires singling out one conformation in a potentially much larger set. A more demanding test for our new similarity indices would be to see if they could be used to gain more insight into the overall distribution of all the structures in the ensemble. A direct way to accomplish this is by using our extended indices as the basis of a new hierarchical agglomerative clustering (HAC) algorithm. The way to do so is relatively simple:

1. We begin by assigning all the objects to separate, disjoint, clusters.
2. If in the *k*^th^ step we have clusters 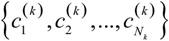, we proceed to the (*k*+1)^th^ iteration by combining the clusters *a* and *b* that give the maximum value of the extended similarity 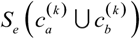.

This is nothing but a natural alternative of the classical HAC algorithms, where we use the *n*-ary indices as linkage criterion. However, there are some important differences with previous methods. For instance, popular single- and complete-linkage criteria measure the relation between clusters in a simple way, using a single point-to-point distance to determine the degree of similarity between clusters. While intuitive, this is a manifestly local approach, in the sense that much of the clusters’ structure is ultimately ignored. On the other hand, by using *n*-ary indices we take into account all the elements at the same time, so this has the potential to provide a better measure of inter-cluster distance.

We tested this hypothesis on two different datasets on protein-DNA binding and protein G folding. For each system, we generate six distinct clusters from MELD simulations and one noisy set to evaluate the robustness of each linkage criterion. Simulation details are shown in Fig.S2 along with pairwise RMSD plots and representative structures for each cluster. The quality of the clustering is obtained using the V-measure, in order to assess both cluster completeness and homogeneity.^26^

As expected, using *n*-ary indices provides a more robust clustering criterion than the local measures. This is particularly evident for the single linkage, where not only we observe the lowest value of the V-measure, but also in system protein G we can see a great dispersion of the possible values of this indicator. Noticeably, the average linkage also results in intermediate-to-poor V-measure values. This is interesting because it shows the limitations of the pairwise similarity metrics, because even when we try to approximate a “global” description by averaging all the pairwise distances, this is still not enough to fully capture the overall relation between the clusters. On the other hand, the *n*-ary indices succeed on this task. We can corroborate this by seeing how the extended RR index consistently provides V values with small dispersion that are close to 1, similarly than those obtained using the Ward criterion. This implies that the effect of maximizing the *n*-ary indices is similar to minimizing the variance of the data. This is in line with previous observations noting that our indices provide a robust description of set compactness. Notice, however, that the good results obtained through Ward linkage are related to the way in which we constructed the dataset, since we used pre-selected clusters with small variance, which is directly related to Ward’s objective function. A more realistic example, using a more realistic dataset with high variance, will be discussed in the next section.

Finally, another crucial advantage of the extended similarity-based HAC is that it gives us a convenient way to estimate the optimal number clusters in the data. The key here is that we can define “cost function”, Δ*_S_*, to evaluate the cost of merging two clusters based on the change in similarity. For example, if we combine clusters *a* and *b* in the *k*^th^ step, the associated cost will be given by:

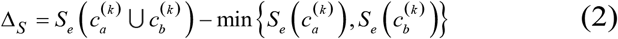

This is a very convenient expression, since we have to calculate the extended similarities anyways, so we do not need to devote any extra computational power to this task (as opposed to other methods to compute the number of clusters, like X-means clustering and information theoretic approaches).^27–29^ Then, whenever we see a steep variation in Δ*_S_* when we go from one iteration to the next. That means that we should not have combined those clusters. We illustrate this behaviour in Fig. 4, where, unsurprisingly, we see that in the last stages of the clustering process the cost increases significantly.

**Figure 3.**
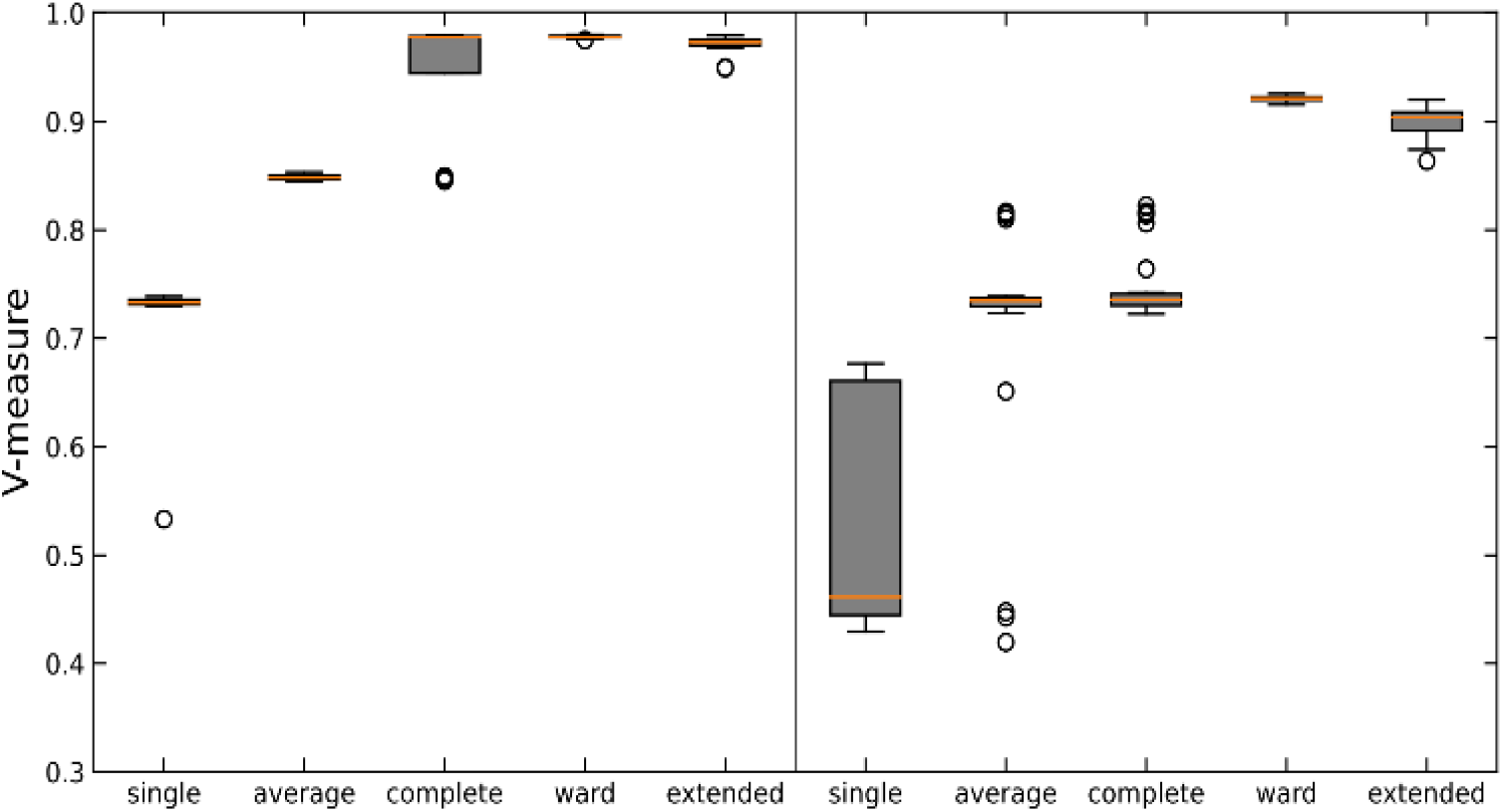
Box plot of V-measure score for different linkages on two synthetic data sets: protein-DNA (left) and protein G (right). Each linkage is tested against 30 subsampled data sets. The orange line in each box represents the median score.

**Figure 4.**
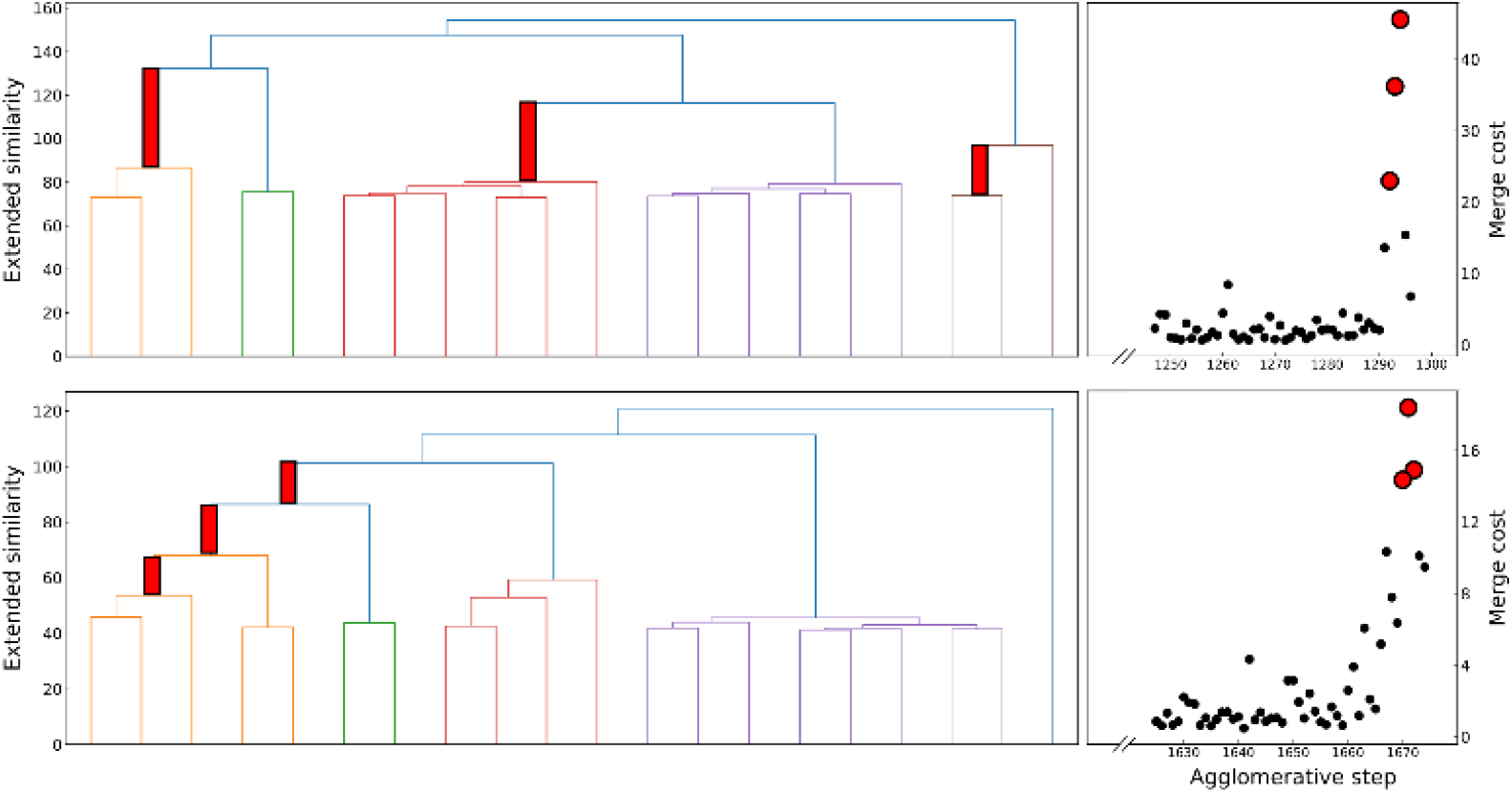
*Left*: hierarchical clustering tree plots on the left show the change of cluster similarity with merging steps for protein-DNA (top) and protein G (down). For clarity, each similarity value is subtracted by the maximum of all similarity values. *Right*: associated merge cost of the last steps.

### Exploring the protein folding landscape

Now we turn our attention to using the extended similarity indices to gain a deeper understanding into the different stages of the protein folding process. To do this, and taking into account the previously discussed properties of our indices, we propose the following workflow:

1. Given a set of structures, find the one that is closest to the reference state. (e.g., by finding the medoid, as detailed above).
2. For all the remaining conformations, calculate their similarity with respect to the reference state.
3. Separate all the structures in sub-sets according to their similarity with respect to the reference state.
4. Cluster the structures in each of the previous sub-sets.

Overall, this novel approach on the analysis of protein folding is extremely efficient. As we have already shown, finding the native structure scales as O(*N*), which is also the scaling of the 2^nd^ step. Finally, notice that we greatly reduce the burden of a traditional clustering algorithm by effectively dividing the data into “bins”, which are then studied individually. In this regard, step 3 effectively serves as a data-reduction technique. The idea of selecting a reduced set of structures (in this case, one) as reference to compare with others in order to describe a larger set is reminiscent of the chemical satellites approach used to visualize the chemical space.^30, 31^ However, in our case, instead of selecting an outlier, we use the medoid of the set (e.g., its most representative point) as reference to “navigate” through the rest of the conformations.

We applied this workflow to the NuG2 dataset (see Fig. 5), where first we selected 5000 random samples from the full dataset, before proceeding through steps 1-4. In this particular case, we separated the data in four sub-sets in the 3^rd^ step: folded (0.06>*csim*), intermediate (0.12>*csim*≥0.06), pre-folded (0.16>*csim*≥0.12), and random coil (*csim*≥0.16), where *csim* stands for the complementary similarity of each conformation. Given the homogeneity of the folded subset and, conversely, the extremely disorganized nature of the random coil conformations, we only performed the clustering on the intermediate and pre-folded states.

**Figure 5.**
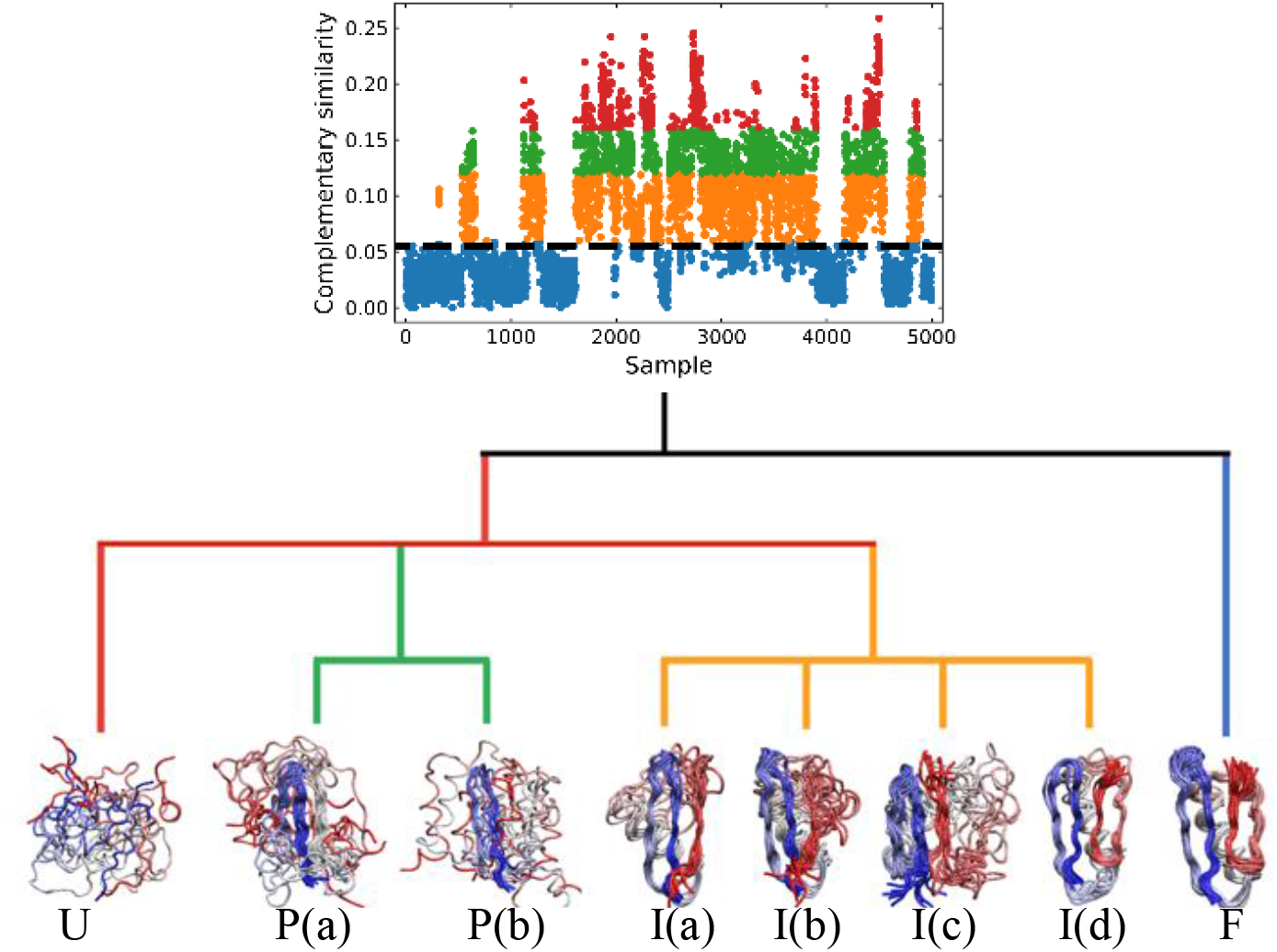
Hierarchical protein folding landscape of NuG2. After calculating complementary similarity and binary similarity (see Fig. S5), the data set is divided into four subsets: folded (blue), intermediate (orange), pre-folded (green) and random coil (red). Hierarchical clustering with extended similarity linkage is performed on pre-folded and intermediate states with representatives in highly populated clusters shown at the bottom.

In the clustering of biological ensemble datasets (and other data science applications), besides the evaluation of the centroid, a stricter assessment of the clustering process is the calculation of the relative population of different clusters. This is particularly relevant for the most populated states, which should manifest the dominating conformations of interest (e.g., in this case, intermediates in the folding process). The clustering using extended similarity indices is also superior to the standard linkage criteria in this regard, when we perform the clustering over the intermediate state datasets (e.g., step 4 in the previous workflow). As shown in Fig. 6, and similarly to the discussion in the previous section, single and average linkage completely failed to identify different clusters in this state and as expected. On the other hand, complete and Ward linkage tend to create one big cluster with high variance. In particular, as the result of its objective function, Ward linkage prefers to make the high-variance cluster more populated, likely due to the penalty to merge with small cluster with little variance (see Fig. S4), which leads to an erroneous representation of the ensemble’s landscape. By comparing with the latest analysis of same NuG2 folding dataset, it is remarkable that all clusters derived here correspond well with essential states along two distinct folding pathways from relaxation mode analysis,^32^ using both structural and time information: after the early formation of N-terminal β-sheet, intermediate states I(a) and I(c) are identified as the most populated clusters before finally folding to native state. In addition, I(d) was identified that corresponds to a near-native state containing a two-residue register shift in the N-terminal hairpin, which was studied further by Schwantes *et al*.^33^ In this way the unfolded clusters have few stabilizing interactions and are internally quite diverse, the folded states are characterized by many stable internal interactions (e.g. hydrogen bonding) leading to a narrow ensemble of structures (lower entropy). In a nutshell, our strategy of pre-classifying all conformations along a density-like similarity coordinate and then performing further clustering with the extended similarity index in regions of interest provides a robust and highly efficient way to analyse challenging biological ensembles.

**Figure 6.**
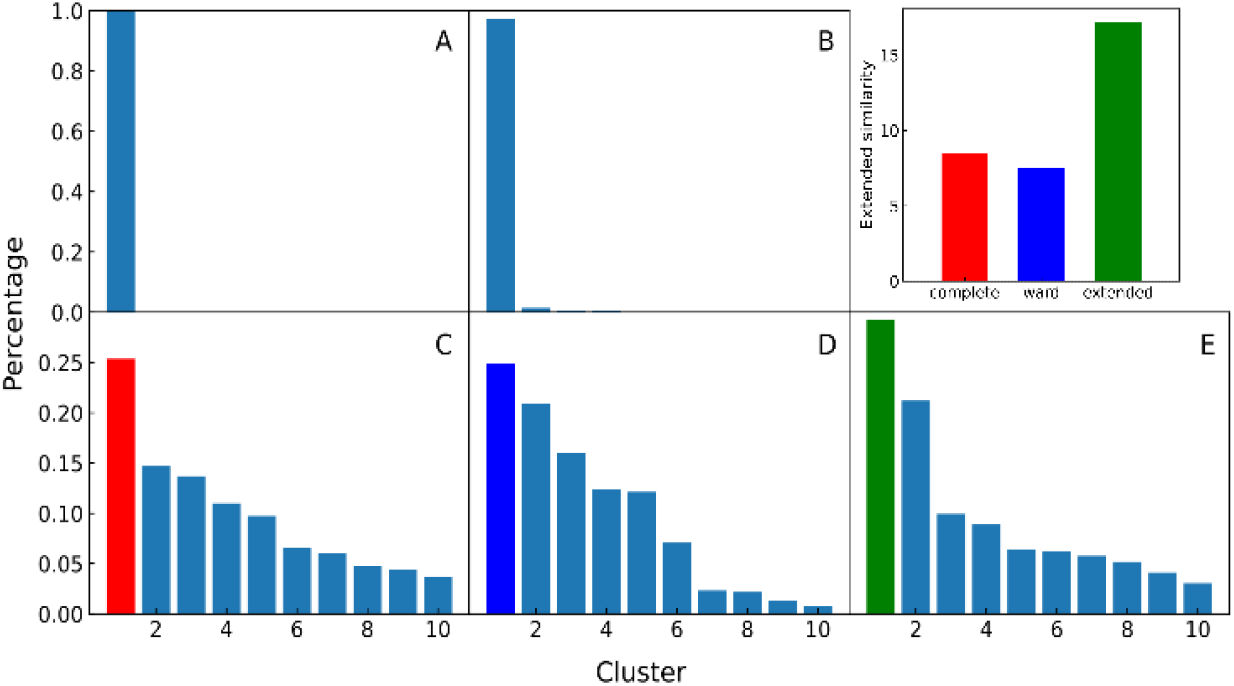
*A-E panels*: Comparison of different linkages in clustering intermediate states. A and B: single linkage and average linkage failed to identify distinct states. C, D and E: complete, Warn and extended linkages. *Top right panel*: Non-weighted Russell-Rao index is calculated for the top cluster of complete, Ward and extended to compare the homogeneity, showing that complete and Ward tend to create a high variance big cluster that leads to erroneous estimate of states population.

## Conclusions

We presented a set of highly efficient analysis tools for biological ensembles based on their inter- or intra-contacts extended similarity. First, the group similarity is defined as a convenient density coordinate to rank a data set, where the highest group similarity effectively corresponds to the medoid (e.g., the ‘most representative element’) of the data set. Unlike other density based clustering algorithms, the group similarity does not require the definition of a hyperparameter, such as a distance cutoff, but nonetheless it scales quadratically with the size of the set.^34^ Additionally, we propose a linear-time algorithm to solve the same medoid problem using extended similarity measures, which follows the insight that removing the most representative point from a cluster will cause the largest decrease of its extended similarity. We also developed a new hierarchical agglomerative clustering method that uses the extended similarity as a linkage criterion. This novel strategy outperforms conventional linkages, which can be explained by the superior description of set similarity provided by our new indices. An additional benefit of our clustering algorithm is the ability to determine the optimal number of clusters for a data set, which we can obtain from a naturally-defined merging cost function defined along the clustering process (and that can be calculated with no extra computational cost). Finally, we combined all these tools to the study of a complex protein folding landscape, taking as test-case the simulation previously performed by Lindorff-Larsen *et al*. of NuG2,^21^ a designed mutant of protein G.^35^ The workflow that we discussed gave excellent results, allowing us to identify the most important states along two different folding pathways. The unprecedented efficiency and versatility of our approach (combining the selection of a convenient reference state, the definition of a density-like coordinate, and a new way of performing the HAC) highlights the potential of the extended similarity measures to gain deeper structural insights from biological ensembles. This could be in turn combined with a time information-based analysis method such as Markov state model to obtain kinetic information.^36^ These results motivate us to explore further generalizations of the extended similarities in the analysis of biological data, particularly, the extension of our formalism in order to work with vectors with continuous components (e.g., so we can directly work with the Cartesian coordinates of a system, without the need to generate the contact maps). Moreover, given the success of the HAC algorithms based on our indices, we are also working on hierarchical divisive clustering algorithms based on extended similarity measures.

## Author Contributions

**L. C.**: Data curation, Formal analysis, Investigation, Methodology, Software, Visualization, Writing-original draft, Writing-review&editing.

**A. P.**: Conceptualization, Funding acquisition, Methodology, Project administration, Resources, Supervision, Writing-original draft, Writing-review&editing.

**R. A. M. Q.**: Conceptualization, Funding acquisition, Methodology, Project administration, Software, Supervision, Writing-original draft, Writing-review&editing.

## Acknowledgements

AP and RAMQ acknowledge support from the University of Florida in the form of startup grants. We thank Emiliano Brini for the protein-protein ensembles as well as D.E Shaw research for their protein folding pathways ensembles.

