## Supplementary Information for "Improving the analysis of biological ensembles through extended similarity measures"

---

[a] Department of Chemistry, University of Florida, Gainesville, FL 32611, USA.
[b] Quantum Theory Project, University of Florida, Gainesville, FL 32611, USA.


**Example calculation of the non-weighted extended Russell-Rao index**

Let us consider the following toy model of binary fingerprints:

$$\begin{matrix} [1 & 0 & 0 & 1 & 1 & 1] \\ [1 & 1 & 1 & 0 & 0 & 1] \\ [1 & 0 & 1 & 1 & 0 & 0] \\ [1 & 0 & 0 & 1 & 1 & 1] \end{matrix} \qquad (0.1)$$

The first step is to calculate the sum of each column:

$$s = [4 \quad 1 \quad 2 \quad 3 \quad 2 \quad 3] \qquad (0.2)$$

Based on these sum values, we decide whether each of the columns can be identified as a similarity counter given by the coincidence of 'on' bits. To do this we need to define a coincidence threshold, $\gamma$, which in this work we take to be equal to:

$$\gamma = n \bmod 2 \qquad (0.3)$$

with *n* being the number of elements that we are comparing. In this case $n = 4$, so $\gamma = 0$.

We then apply the following criterion: every column such that its sum, $s_k$, satisfies the inequality:

$$2s_k - n > \gamma \qquad (0.4)$$

will correspond to a '1-similarity'. In this case, the 1st, 4th, and 6th columns will correspond to 1-similarities.

Finally, the extended non-weighted Russell-Rao index, $S_{eRR}$, will be given by:

$$S_{eRR} = \frac{\sum_{1-sim} \frac{2s_k - n}{n}}{p} \qquad (0.5)$$

here, *p* represents the length of the vectors being compared, and notice that the summation only runs over the 1-similarity columns. That is, for the data presented in Eq. (0.1):

$$S_{eRR} = \frac{1 + \frac{1}{2} + \frac{1}{2}}{6} = \frac{1}{3} \qquad (0.6)$$

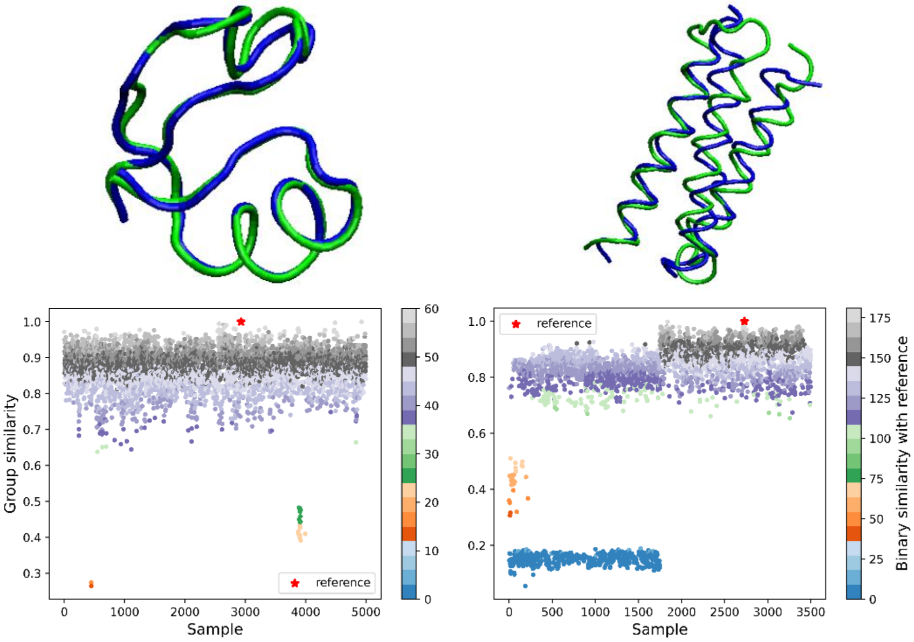


Figure S1. Top plots show the conformation with maximum group similarity for NTL9 (left) and A3D (right) in blue aligned to experimental structure in green. Bottom plots show the group similarity of all samples for ASTP-7041 (left) and 1NS1 (right).

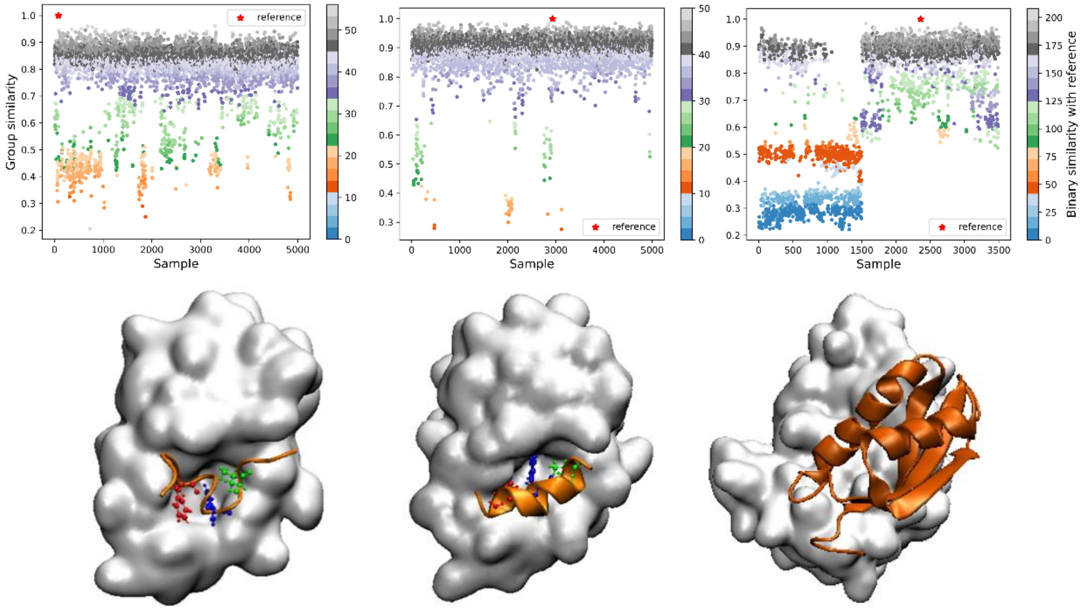


Figure S2. Additional tests for p53 (left), pdiq (middle) and 2MMA (right). Group similarity plots are shown on top and the bottom indicates the representative structures for each data set.

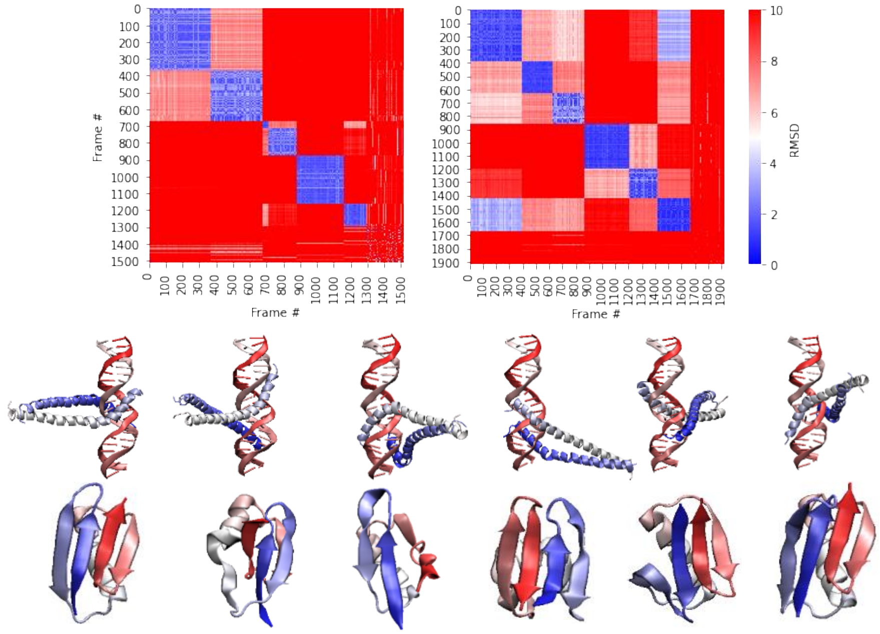


Figure S3**.** Pairwise RMSD plot of generated samples are shown on top for protein-DNA system (left) and protein G (right). Representatives of each cluster are shown at bottom. For each system, we picked six distinct seeds from MELD ensembles generated by selectively enforcing either distance restraints between protein and DNA or secondary structure restraints on alpha helix and two hairpins of protein G, then we choose the most similar conformations against each seed based on RMSD from the rest of ensemble. In addition, we added one noisy set to evaluate the robustness of each linkage criterion.

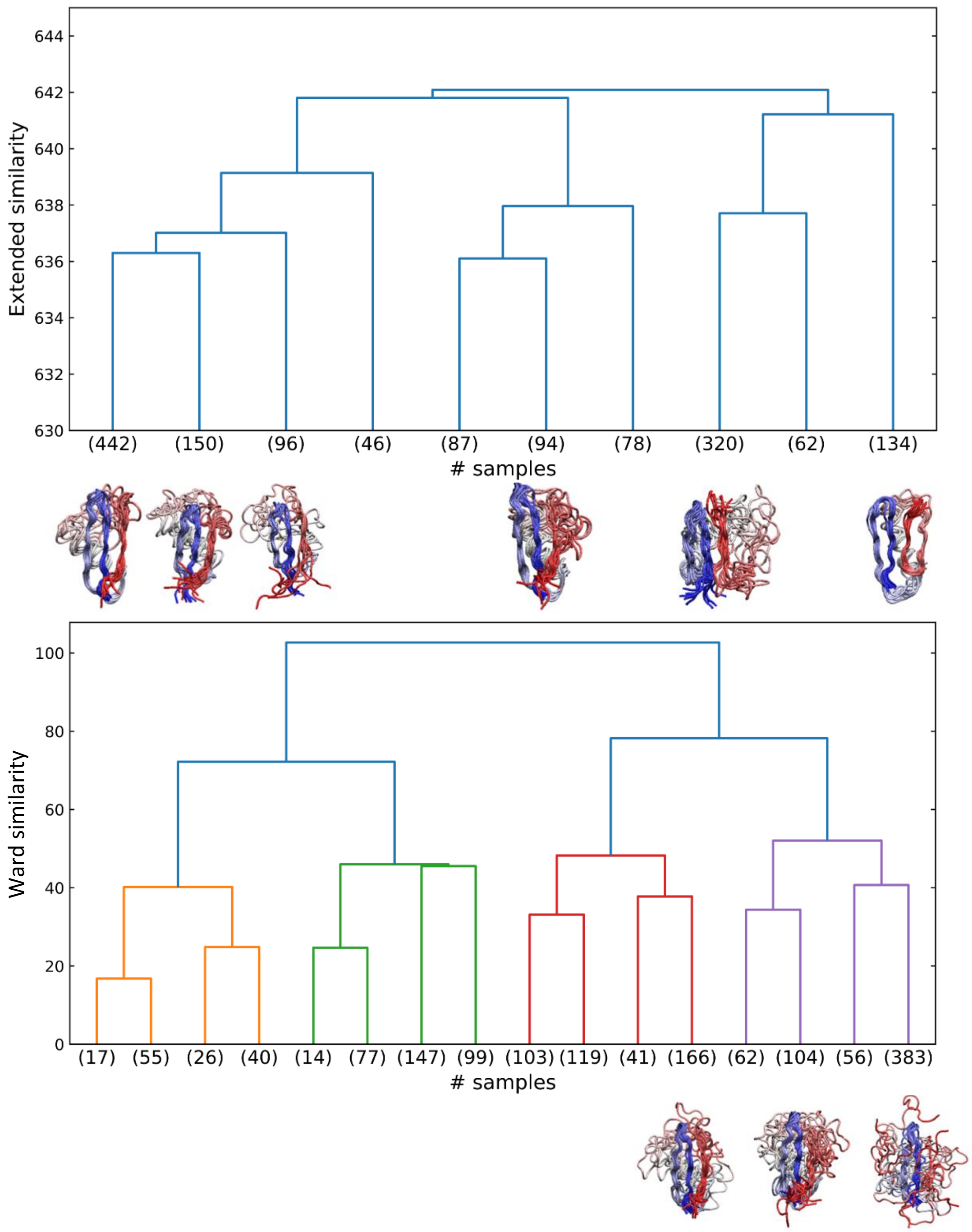


Figure S4. Hierarchical clustering result on intermediate state and representative samples for high population states by using extended similarity (top) and Ward linkage (bottom). The extended similarity value shown is subtracted by the maximum of all similarity values. The bottom plot shows the highest populated cluster using Ward linkage has high variance with many dissimilar conformations.

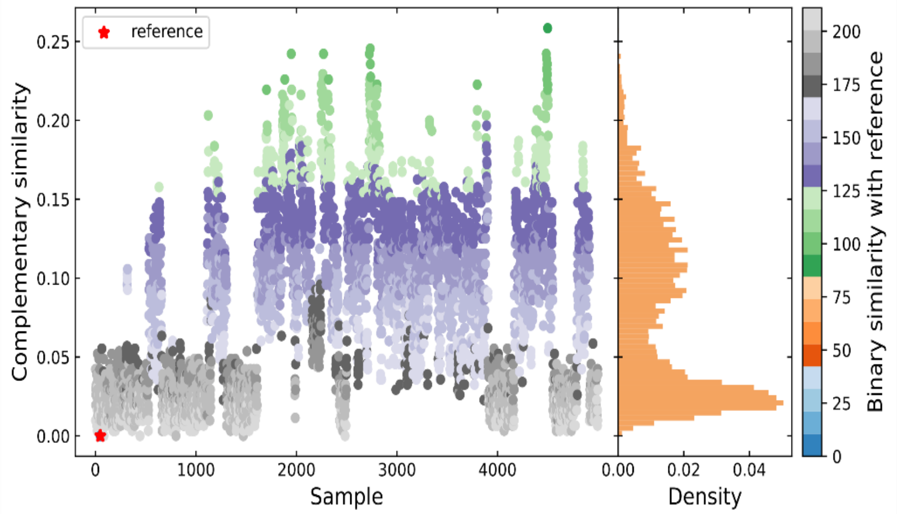


Figure S5. Selection strategy for NuG2 folding ensemble analysis. The color in the scatter plot represents binary similarity (the coincidence value of 1 in both fingerprints) of each sample with the reference conformation, which corresponds to the minimum of the complementary similarity (e.g., the medoid).
